# Large-scale cognitive GWAS Meta-Analysis Reveals Tissue-Specific Neural Expression and Potential Nootropic Drug Targets

**DOI:** 10.1101/176842

**Authors:** Max Lam, Joey W. Trampush, Jin Yu, Emma Knowles, Gail Davies, David C. Liewald, John M. Starr, Srdjan Djurovic, Ingrid Melle, Kjetil Sundet, Andrea Christoforou, Ivar Reinvang, Pamela DeRosse, Astri J. Lundervold, Vidar M. Steen, Thomas Espeseth, Katri Räikkönen, Elisabeth Widen, Aarno Palotie, Johan G. Eriksson, Ina Giegling, Bettina Konte, Panos Roussos, Stella Giakoumaki, Katherine E. Burdick, Antony Payton, William Ollier, Ornit Chiba-Falek, Deborah K. Attix, Anna C. Need, Elizabeth T. Cirulli, Aristotle N. Voineskos, Nikos C. Stefanis, Dimitrios Avramopoulos, Alex Hatzimanolis, Dan E. Arking, Nikolaos Smyrnis, Robert M. Bilder, Nelson A. Freimer, Tyrone D. Cannon, Edythe London, Russell A. Poldrack, Fred W. Sabb, Eliza Congdon, Emily Drabant Conley, Matthew A. Scult, Dwight Dickinson, Richard E. Straub, Gary Donohoe, Derek Morris, Aiden Corvin, Michael Gill, Ahmad R. Hariri, Daniel R. Weinberger, Neil Pendleton, Panos Bitsios, Dan Rujescu, Jari Lahti, Stephanie Le Hellard, Matthew C. Keller, Ole A. Andreassen, Ian J. Deary, David C. Glahn, Anil K. Malhotra, Todd Lencz

**Affiliations:** Institute of Mental Health, Singapore; BrainWorkup, LLC, Los Angeles, CA; Division of Psychiatry Research, The Zucker Hillside Hospital, Glen Oaks, NY, USA; Department of Psychiatry, Yale University School of Medicine, New Haven, CT, USA; Centre for Cognitive Ageing and Cognitive Epidemiology, University of Edinburgh, Edinburgh, United Kingdom; Department of Psychology, University of Edinburgh, Edinburgh, United Kingdom; Alzheimer Scotland Dementia Research Centre, University of Edinburgh, Edinburgh, United Kingdom; Department of Medical Genetics, Oslo University Hospital, University of Bergen, Oslo, Norway; NORMENT, K.G. Jebsen Centre for Psychosis Research, University of Bergen, Bergen, Norway; Division of Mental Health and Addiction, Oslo University Hospital, Oslo, Norway; Department of Psychology, University of Oslo, Oslo, Norway; Dr. Einar Martens Research Group for Biological Psychiatry, Center for Medical Genetics and Molecular Medicine, Haukeland University Hospital, Bergen, Norway; Department of Biological and Medical Psychology, University of Bergen, Norway; Institute of Behavioural Sciences, University of Helsinki, Helsinki, Finland; Institute for Molecular Medicine Finland (FIMM), University of Helsinki, Finland; Wellcome Trust Sanger Institute, Wellcome Trust Genome Campus, Cambridge, United Kingdom; Department of Medical Genetics, University of Helsinki and University Central Hospital, Helsinki, Finland; Department of General Practice, University of Helsinki and Helsinki University Hospital, Helsinki, Finland; National Institute for Health and Welfare, Helsinki, Finland; Folkhälsan Research Center, Helsinki, Finland; Department of Psychiatry, Martin Luther University of Halle-Wittenberg, Halle, Germany; Department of Psychiatry, Icahn School of Medicine at Mount Sinai, New York, NY, USA; Department of Genetics and Genomic Science and Institute for Multiscale Biology, Icahn School of Medicine at Mount Sinai, New York, NY, USA; Mental Illness Research, Education, and Clinical Center (VISN 2), James J. Peters VA Medical Center, Bronx, NY, USA; Department of Psychology, University of Crete, Greece; Department of Psychiatry - Brigham and Women’s Hospital; Harvard Medical School; Boston MA; Centre for Epidemiology, Division of Population Health, Health Services Research & Primary Care, The University of Manchester, Manchester, United Kingdom; Centre for Integrated Genomic Medical Research, Institute of Population Health, University of Manchester, Manchester, United Kingdom; Department of Neurology, Bryan Alzheimer’s Disease Research Center, and Center for Genomic and Computational Biology, Duke University Medical Center, Durham, NC, USA; Psychiatry and Behavioral Sciences, Division of Medical Psychology, and Department of Neurology, Duke University Medical Center, Durham, NC, USA; Division of Brain Sciences, Department of Medicine, Imperial College, London, UK; Human Longevity Inc, Durham, NC, USA; Campbell Family Mental Health Institute, Centre for Addiction and Mental Health, University of Toronto, Toronto, Canada; Department of Psychiatry, National and Kapodistrian University of Athens Medical School, Eginition Hospital, Athens, Greece; University Mental Health Research Institute, Athens, Greece; Neurobiology Research Institute, Theodor-Theohari Cozzika Foundation, Athens, Greece; Department of Psychiatry, Johns Hopkins University School of Medicine, MD, Baltimore, USA; McKusick-Nathans Institute of Genetic Medicine, Johns Hopkins University School of Medicine, MD, Baltimore, USA; UCLA Semel Institute for Neuroscience and Human Behavior, Los Angeles, CA, USA; Department of Psychology, Yale University, New Haven, CT, USA; Department of Psychology, Stanford University, Palo Alto, CA, USA; Robert and Beverly Lewis Center for Neuroimaging, University of Oregon, Eugene, OR, USA; 23andMe, Inc., Mountain View, CA, USA; Laboratory of NeuroGenetics, Department of Psychology & Neuroscience, Duke University, Durham, NC, USA; Clinical and Translational Neuroscience Branch, Intramural Research Program, National Institute of Mental Health, National Institute of Health, Bethesda, MD, USA; Lieber Institute for Brain Development, Johns Hopkins University Medical Campus, Baltimore, MD, USA; Neuroimaging, Cognition & Genomics (NICOG) Centre, School of Psychology and Discipline of Biochemistry, National University of Ireland, Galway, Ireland; Neuropsychiatric Genetics Research Group, Department of Psychiatry and Trinity College Institute of Neuroscience, Trinity College Dublin, Dublin, Ireland; Division of Neuroscience and Experimental Psychology/ School of Biological Sciences, Faculty of Biology Medicine and Health, University of Manchester, Manchester Academic Health Science Centre, Salford Royal NHS Foundation Trust, Manchester, United Kingdom; Department of Psychiatry and Behavioral Sciences, Faculty of Medicine, University of Crete, Heraklion, Crete, Greece; Helsinki Collegium for Advanced Studies, University of Helsinki, Helsinki, Finland; Institute for Behavioral Genetics, University of Colorado, Boulder, Colorado; Institute of Clinical Medicine, University of Oslo, Oslo, Norway; Department of Psychiatry, Hofstra Northwell School of Medicine, Hempstead, New York; Center for Psychiatric Neuroscience, Feinstein Institute for Medical Research, Manhasset New York

## Abstract

Neurocognitive ability is a fundamental readout of brain function, and cognitive deficits are a critical component of neuropsychiatric disorders, yet neurocognition is poorly understood at the molecular level. In the present report, we present the largest genome-wide association studies (GWAS) of cognitive ability to date (N=107,207), and further enhance signal by combining results with a large-scale GWAS of educational attainment. We identified 70 independent genomic loci associated with cognitive ability, 34 of which were novel. A total of 350 genes were implicated, and this list showed significant enrichment for genes associated with Mendelian disorders with an intellectual disability phenotype. Competitive pathway analysis of gene results implicated the biological process of neurogenesis, as well as the gene targets of two pharmacologic agents: cinnarizine, a T-type calcium channel blocker; and LY97241, a potassium channel inhibitor. Transcriptome-wide analysis revealed that the implicated genes were strongly expressed in neurons, but not astrocytes or oligodendrocytes, and were more strongly associated with fetal brain expression than adult brain expression. Several tissue-specific gene expression relationships to cognitive ability were observed (for example, *DAG1* levels in the hippocampus). Finally, we report novel genetic correlations between cognitive ability and disparate phenotypes such as maternal age at first birth and number of children, as well as several autoimmune disorders.

## Introduction

Genome-wide association studies (GWAS) have been highly successful at uncovering hundreds of genetic loci associated with heritable quantitative traits such as height^1^ and weight^2^ (body mass index). However, identifying genetic loci underlying cognitive ability has been much more challenging, despite comparably high levels of heritability as determined by both classical twin studies^3^ and molecular genetic studies^4^. Uncovering the molecular genetic basis of individual differences in cognitive performance can have a significant impact on our understanding of neuropsychiatric disorders, which are both phenotypically^5-8^ and genetically^9-11^ correlated with cognition, as well as numerous non-psychiatric health-relevant phenotypes^12^ that also are significantly genetically correlated with cognitive function.

In part, the difficulty with cognitive GWAS may be caused by the relative degree of heterogeneity in the measurement of the cognitive phenotype. Traditionally, general cognitive ability (*g*) has been defined as a latent trait underlying shared variance across multiple subdomains of cognitive performance, psychometrically obtained as the first principal component of several distinct neuropsychological test scores^13^. Using this approach, several cognitive GWAS with fewer than 20,000 subjects yielded no genome-wide significant (GWS) effects^4,9,14^, while a few GWS loci were identified in large GWAS of 35,298^15^ subjects and 53,949^16^ subjects, respectively. Notably, these efforts involved meta-analysis across cohorts using different sets of cognitive tests to derive the principal component score, which may have reduced power. By contrast, two independent GWAS of height with sample sizes of approximately 30,000 subjects each yielded 20-30 GWS hits^17,18^; allelic effect sizes were ∼2-5 times larger than the largest obtained in cognitive GWAS^19^.

Given the small effect sizes observed in cognitive GWAS, it has become evident that greatly increased sample sizes will be required to ascend the GWAS yield curve. Very recently, a cognitive GWAS^20^ was able to leverage a very brief (two-minute) measure of fluid intelligence, highly correlated with psychometrically defined *g*, obtained in over 50,000 subjects. In combination with several traditional cognitive GWAS cohorts, total sample size was 78,308. This sample size permitted discovery of 18 independent GWS allelic loci, as well as numerous additional loci from gene-based analysis. This report was critical in demonstrating that signal could be enhanced by combining data from cohorts with brief measures of intelligence with more traditional cognitive GWAS.

Yet another approach to enhancing power in cognitive GWAS has focused on educational attainment as a proxy phenotype^21^. It is acknowledged that this phenotype is ‘noisy,’ as it is influenced by non-cognitive genetic^22^ (e.g., personality) and environmental^23^ (e.g., socio-economic) factors; consequently, observed allelic effect sizes have been even smaller than those obtained for GWAS of *g*^24^. However, by utilizing a single-item measure (years of education completed), obtained incidentally in large studies of other phenotypes, this approach has allowed investigators to obtain extremely large sample sizes. A recent study of educational attainment in nearly 300,000 individuals identified 74 independent GWS loci^25^. Notably, the genetic correlation between educational attainment and psychometric *g* is very high, consistently reported in the range of .70-.75^15,16,20,25,26^.

Thus, cognitive GWAS can be further enhanced by combining information from these large studies of educational attainment with studies of test-based cognitive performance. A new technique called multi-trait analysis of GWAS (MTAG)^27^ has been developed which permits integration of GWAS data across related traits, accounting for the possibility of overlapping samples across studies, and requiring only summary statistics. Notably, the developers of MTAG demonstrated its accuracy and utility in a study of traits that also demonstrate genetic correlations in the range of ∼.70-.75 (depression, neuroticism, and subjective well-being). MTAG is able to quantify the degree of “boost” to the signal of a single-trait GWAS, providing an estimate of observed sample size, and providing summary statistics (allelic weights) that can then be utilized in all downstream annotation pipelines available for GWAS output.

In the present study, we first utilized GWAS meta-analysis to combine our prior COGENT GWAS^15^ of psychometrically defined *g* with the recently reported GWAS^20^ relying primarily on the brief measure, resulting in a combined cohort of N=107,207 non-overlapping samples. Next, we utilized MTAG to combine these results with the large-scale GWAS of educational attainment, resulting in enhanced power. At each step, we performed both allelic and gene-based tests. We then performed downstream analyses on the resulting MTAG summary statistics, including: 1) competitive gene set analyses to identify key biological processes and potential drug targets implicated; 2) stratified linkage disequilibrium score regression (LDSC) to identify differential cell type expression; 3) transcriptome-wide association study (TWAS) methods, to identify specific effects of altered gene expression in the brain on cognition; and 4) LDSC to identify genetic correlations with other anthropometric and biomedical phenotypes.

## Results

### Fixed Effect Meta-Analysis: Cognitive Performance GWAS

Fixed effect meta-analysis of all non-overlapping cohorts from the two GWAS of cognitive performance (total N = 107,207) identified 28 independent genomic loci reaching genome-wide significance (GWS, p<5E-08), using default clumping parameters from the FUMA^28^ pipeline (Figure 1a); this represents a 55.6% increase in loci compared to the previous GWAS^20^ of cognitive performance. Two of these loci each contained two uncorrelated variants with independent effects, resulting in 30 independent lead SNPs. As demonstrated in the QQ plot (Supplementary Figure 1), statistical inflation was quite modest for a large study of a highly polygenic trait (λ=1.23; λ_1000_=1.001; LD score intercept=1.03), and overall SNP heritability was .168. Of the 28 GWS loci, 12 are novel and not previously reported as GWS in published studies of cognitive or educational phenotypes (Supplementary Table 2). The majority of the 5,610 markers reaching a nominal significance threshold were intronic SNPs followed by those in the intergenic regions (Supplementary Table 3 and Supplementary Figure 2). As shown in Supplementary Table 4, several of the GWS loci overlap with loci related to schizophrenia, bipolar disorder, and other neuropsychiatric phenotypes, as well as obesity/body mass index and other traits.

**Figure 1.**
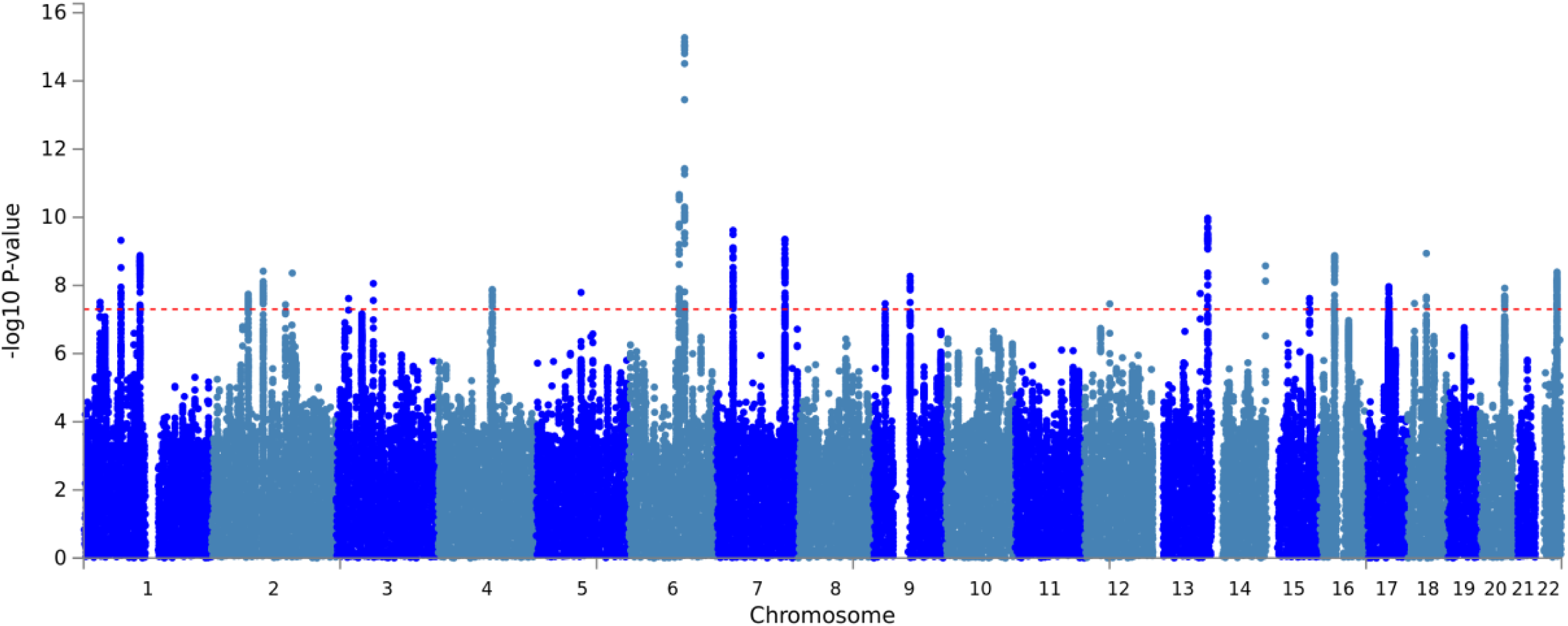
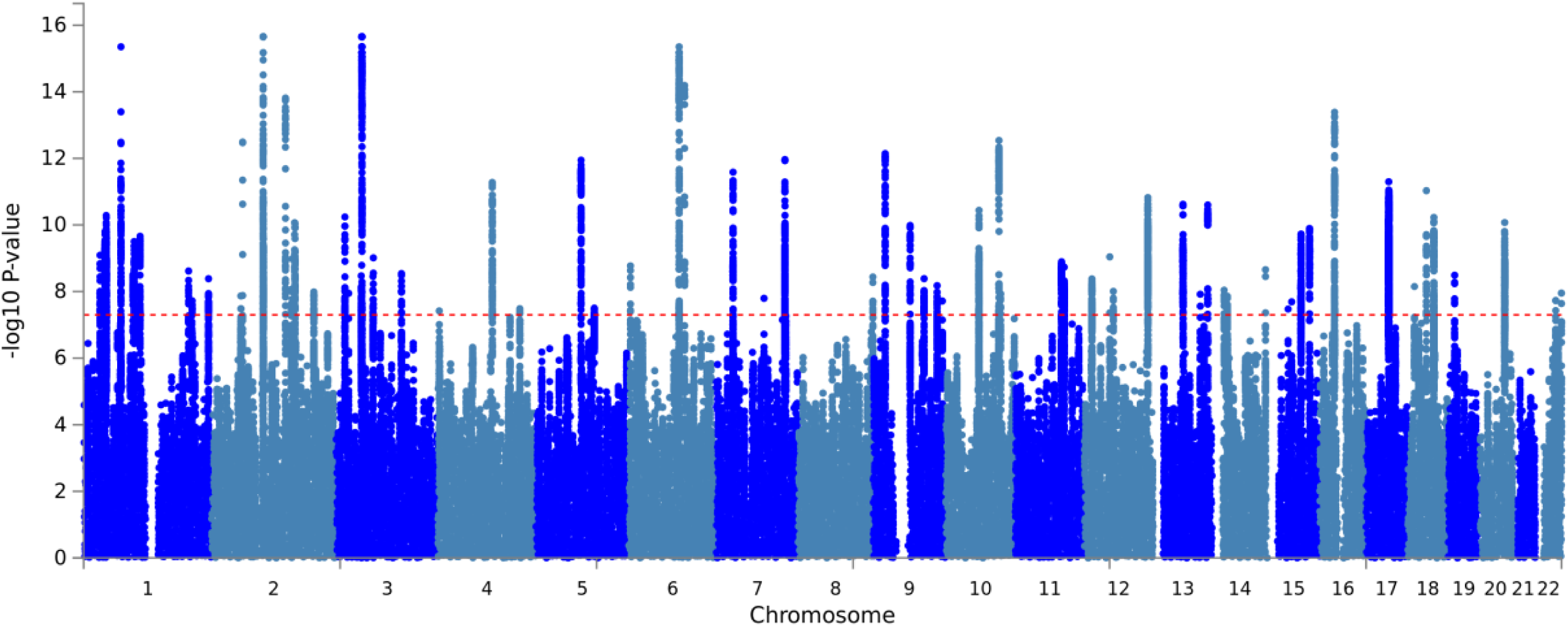
Manhattan plot depicting results of GWAS meta-analysis for cognitive performance (Figure 1a) and MTAG of cognitive performance with educational attainment. Dotted red line indicates threshold for genome-wide significance (P< 5x10^-8^).

**Figure 2.**
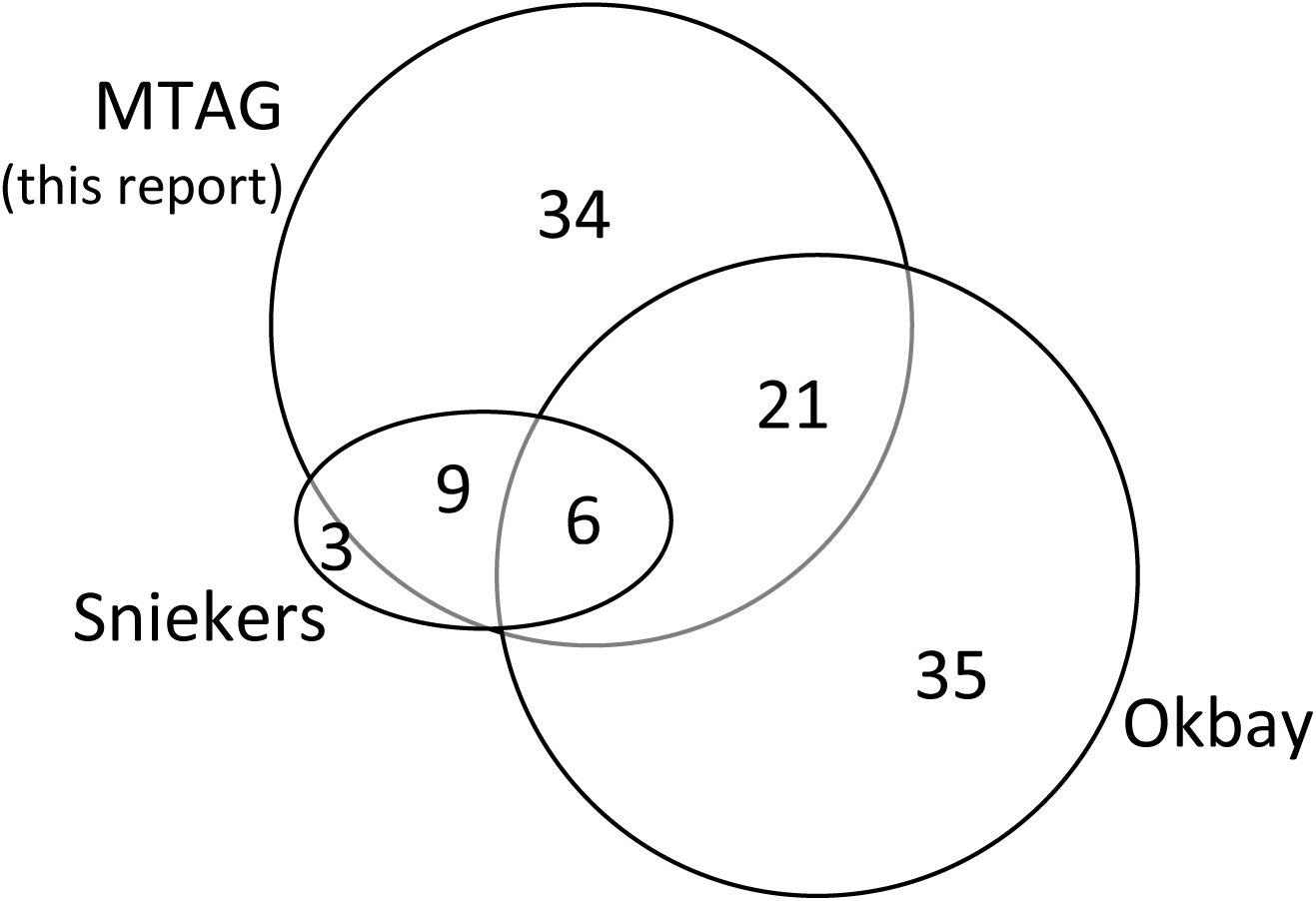
Proportional Venn diagram depicting overlap and independence of genome-wide significant SNP loci observed in three studies: the MTAG analysis of the present report; the cognitive performance GWAS reported by Sniekers et al.^20^; and the educational attainment GWAS of Okbay et al.^25^.

The significant loci harbored 88 known protein coding genes (Supplementary Table 5), about half of which were in three large regions (Supplementary Figure 3), including two well-characterized regions: the distal 16p11.2 region, in which deletions have been associated with schizophrenia and other neuropsychiatric phenotypes^29^, and the 17q21 region, in which inversions have been associated with neuropsychiatric disorders^30,31^. Using MAGMA^32^ gene-based tests, 73 genes were genome-wide significant (Supplementary Figure 4; Supplementary Table 6), of which 39 were overlapping with the 88 genes noted above, resulting in a total of 122 genes with GWS evidence of association to cognitive performance.

**Figure 3.**
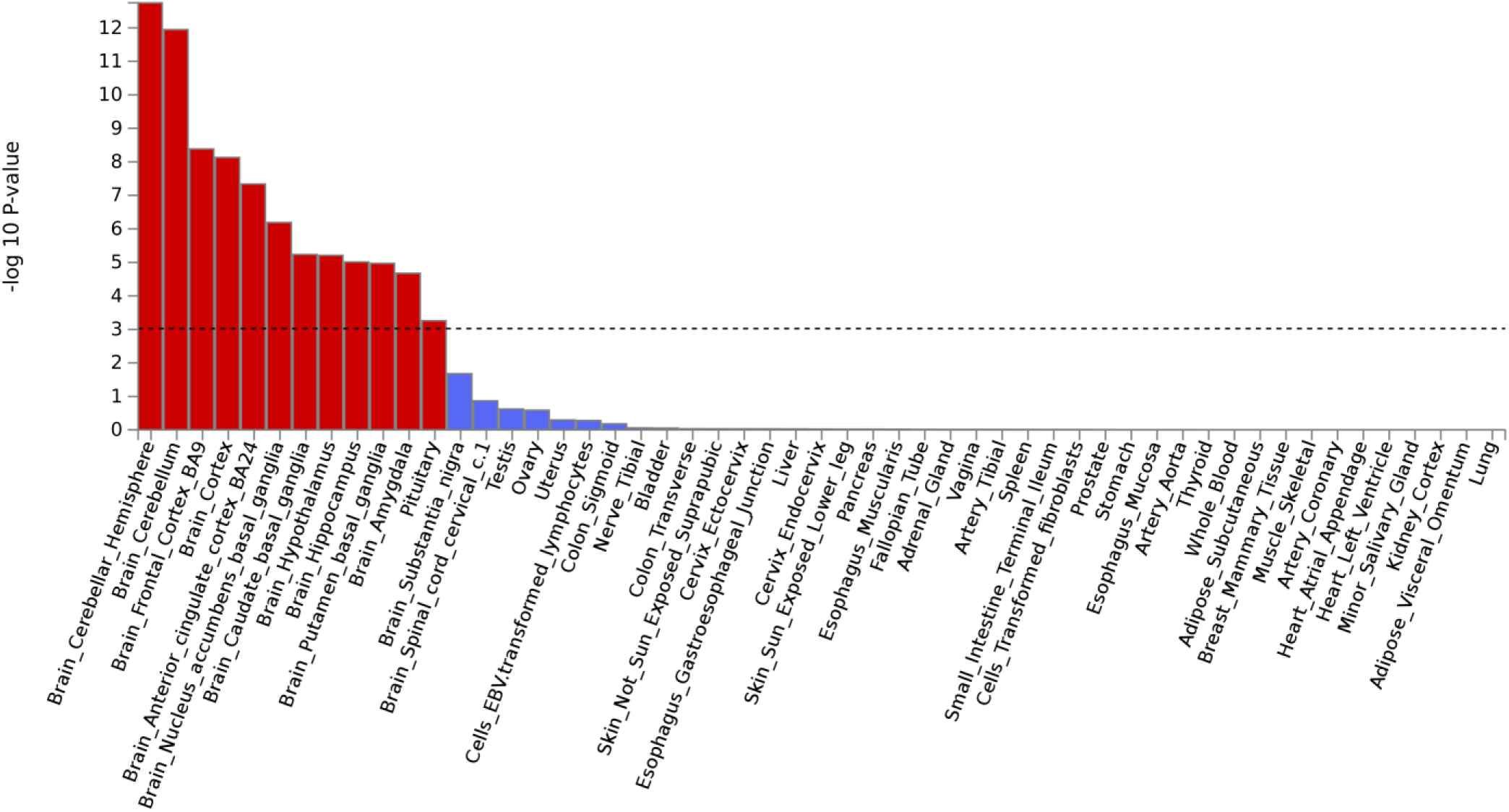
Tissue expression profile analysis for genome-wide significant genes (as defined by MAGMA) emerging from the MTAG analysis. Gene results were significantly enriched for expression in nearly all central nervous system tissues (except for substantia nigra and spinal cord), but no tissues outside the CNS.

### MTAG: Combining Cognitive Performance and Educational Attainment GWAS

MTAG analysis combining the cognitive performance results obtained above with the large education attainment GWAS previously reported^25^, resulted in a 75% enrichment of statistical power, effectively boosting the original sample size of N = 107,207 to a GWAS equivalent of N = 187,812 (Table 1). Default clumping procedures revealed that 70 independent genomic loci reached genome-wide significance, with 82 independent SNPs (Figure 1b). Similar to the GWAS results above, the QQ plot (Supplementary Figure 5), demonstrated polygenicity without substantial statistical inflation (λ=1.28; λ_1000_=1.001; LD score intercept=0.91), and overall SNP heritability was 0.336. Of the 70 GWS loci, 34 are novel and not previously reported as GWS in published studies of cognitive or educational phenotypes (Figure 2; Supplementary Table 7). The majority of the 13,549 SNPs reaching a nominal significance threshold were intergenic or intronic (Supplementary Table 8; Supplementary Figures 6 & 7). As is typically the case in GWAS, few significant SNPs were exonic; GWS variants causing protein-coding changes rated as damaging by either Polyphen or SIFT are listed in Supplementary Table 9. Variants in only four genes demonstrated converging evidence for being damaging to protein structure: three at chromosome 17q21.31 (*KANSL1*, *MAPT*, and *SPPL2C*), as well as one at chromosome 3p21.31 (*MST1*). GWAS catalog annotations are listed in Supplementary Table 10.

**Table 1.** MTAG model output

| Trait | N<br>(max) | No. of<br>SNPs | GWAS<br>mean<br>$\chi^2$ | MTAG<br>mean<br>$\chi^2$ | GWAS<br>Equivalent<br>N | MTAG<br>Signal<br>Boost |
| --- | --- | --- | --- | --- | --- | --- |
| Cognition + Intelligence | 107207 | 7333852 | 1.245 | 1.429 | 187812 | 75% |
| Education | 328917 | 7333852 | 1.638 | 1.693 | 357156 | 8.6% |

Within the GWS loci, 267 protein coding genes were identified (Supplementary Table 11). Additionally, 257 genes were significant in MAGMA gene-based tests (Supplementary Figure 8; Supplementary Table 12); of these, 83 genes were non-overlapping with the 267 genes with SNP GWS loci, resulting in a total of 350 genes receiving GWS support from the MTAG results. We compared this list of 350 genes with a list of 621 genes known to cause autosomal dominant or autosomal recessive Mendelian disorders featuring intellectual disability^33,34^. As shown in Table 2, a total of 23 genes identified by MTAG appeared on this list, representing a 2-fold enrichment over chance (p=0.001).

**Table 2.**
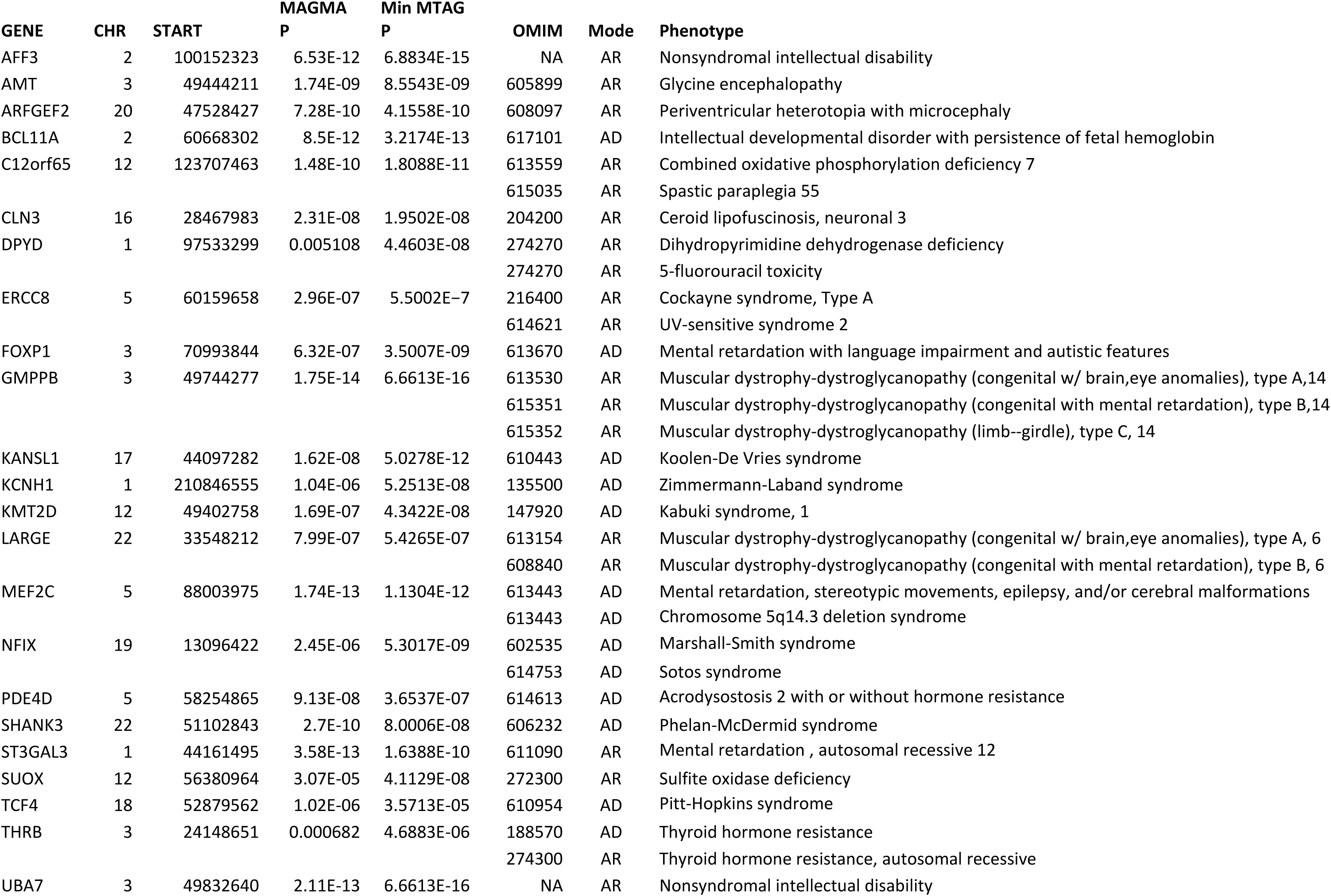

| GENE | CHR | START | MAGMA<br>P | Min MTAG<br>P | OMIM | Mode | Phenotype |
| --- | --- | --- | --- | --- | --- | --- | --- |
| AFF3 | 2 | 100152323 | 6.53E-12 | 6.8834E-15 | NA | AR | Nonsyndromal intellectual disability |
| AMT | 3 | 49444211 | 1.74E-09 | 8.5543E-09 | 605899 | AR | Glycine encephalopathy |
| ARFGEF2 | 20 | 47528427 | 7.28E-10 | 4.1558E-10 | 608097 | AR | Periventricular heterotopia with microcephaly |
| BCL11A | 2 | 60668302 | 8.5E-12 | 3.2174E-13 | 617101 | AD | Intellectual developmental disorder with persistence of fetal hemoglobin |
| C12orf65 | 12 | 123707463 | 1.48E-10 | 1.8088E-11 | 613559 | AR | Combined oxidative phosphorylation deficiency 7 |
|  |  |  |  |  | 615035 | AR | Spastic paraplegia 55 |
| CLN3 | 16 | 28467983 | 2.31E-08 | 1.9502E-08 | 204200 | AR | Ceroid lipofuscinosis, neuronal 3 |
| DPYD | 1 | 97533299 | 0.005108 | 4.4603E-08 | 274270 | AR | Dihydropyrimidine dehydrogenase deficiency |
|  |  |  |  |  | 274270 | AR | 5-fluorouracil toxicity |
| ERCC8 | 5 | 60159658 | 2.96E-07 | 5.5002E-7 | 216400 | AR | Cockayne syndrome, Type A |
|  |  |  |  |  | 614621 | AR | UV-sensitive syndrome 2 |
| FOXP1 | 3 | 70993844 | 6.32E-07 | 3.5007E-09 | 613670 | AD | Mental retardation with language impairment and autistic features |
| GMPPB | 3 | 49744277 | 1.75E-14 | 6.6613E-16 | 613530 | AR | Muscular dystrophy-dystroglycanopathy (congenital w/ brain,eye anomalies), type A,14 |
|  |  |  |  |  | 615351 | AR | Muscular dystrophy-dystroglycanopathy (congenital with mental retardation), type B,14 |
|  |  |  |  |  | 615352 | AR | Muscular dystrophy-dystroglycanopathy (limb--girdle), type C, 14 |
| KANSL1 | 17 | 44097282 | 1.62E-08 | 5.0278E-12 | 610443 | AD | Koolen-De Vries syndrome |
| KCNH1 | 1 | 210846555 | 1.04E-06 | 5.2513E-08 | 135500 | AD | Zimmermann-Laband syndrome |
| KMT2D | 12 | 49402758 | 1.69E-07 | 4.3422E-08 | 147920 | AD | Kabuki syndrome, 1 |
| LARGE | 22 | 33548212 | 7.99E-07 | 5.4265E-07 | 613154 | AR | Muscular dystrophy-dystroglycanopathy (congenital w/ brain,eye anomalies), type A, 6 |
|  |  |  |  |  | 608840 | AR | Muscular dystrophy-dystroglycanopathy (congenital with mental retardation), type B, 6 |
| MEF2C | 5 | 88003975 | 1.74E-13 | 1.1304E-12 | 613443 | AD | Mental retardation, stereotypic movements, epilepsy, and/or cerebral malformations |
|  |  |  |  |  | 613443 | AD | Chromosome 5q14.3 deletion syndrome |
| NFIX | 19 | 13096422 | 2.45E-06 | 5.3017E-09 | 602535 | AD | Marshall-Smith syndrome |
|  |  |  |  |  | 614753 | AD | Sotos syndrome |
| PDE4D | 5 | 58254865 | 9.13E-08 | 3.6537E-07 | 614613 | AD | Acrodysostosis 2 with or without hormone resistance |
| SHANK3 | 22 | 51102843 | 2.7E-10 | 8.0006E-08 | 606232 | AD | Phelan-McDermid syndrome |
| ST3GAL3 | 1 | 44161495 | 3.58E-13 | 1.6388E-10 | 611090 | AR | Mental retardation , autosomal recessive 12 |
| SUOX | 12 | 56380964 | 3.07E-05 | 4.1129E-08 | 272300 | AR | Sulfite oxidase deficiency |
| TCF4 | 18 | 52879562 | 1.02E-06 | 3.5713E-05 | 610954 | AD | Pitt-Hopkins syndrome |
| THRB | 3 | 24148651 | 0.000682 | 4.6883E-06 | 188570 | AD | Thyroid hormone resistance |
|  |  |  |  |  | 274300 | AR | Thyroid hormone resistance, autosomal recessive |
| UBA7 | 3 | 49832640 | 2.11E-13 | 6.6613E-16 | NA | AR | Nonsyndromal intellectual disability |

As a formal validation that the MTAG methodology successfully predicts phenotype variance for cognitive performance, MTAG was re-analyzed, excluding the ASPIS and GCAP datasets from the COGENT cohorts; these datasets were held out as target cohorts used for calculation of polygenic risk score modelling for “g”. Despite the relatively small size of these hold-out cohorts, results show strongly significant polygenic prediction of “g” using MTAG-derived allele weights (Supplementary Figures 9 and 10), accounting for more than 4% of the variance in the GCAP cohort. For both cohorts, polygenic prediction began to drop at P_T_ thresholds above 0.05, suggesting that there may be some degree of saturation of signal beyond the nominal 0.05 significance level.

### Gene Expression and Competitive Pathway Analysis

Downstream MAGMA expression profiles and competitive pathway analysis were conducted as part of the FUMA pipeline. MAGMA tissue expression profile analysis revealed that genes emerging from the MTAG analysis were significantly enriched for expression in nearly all central nervous system tissues (except for substantia nigra and spinal cord), and that this enrichment was exclusive to neural tissues (Figure 3). Notably, the strongest enrichment was observed for genes expressed in the cerebellum, followed by cortex, and slightly weaker (but still strongly significant) enrichment in subcortical and limbic structures.

Competitive pathway analysis (based on gene ontology categories) for GWS MAGMA genes identified by MTAG revealed significant enrichment of neuronal and synaptic cellular components, as well as the biological process of neurogenesis (Table 3a). Competitive pathway analysis for drug pathways^35^ revealed that two drugs were significantly associated with the MTAG results (Table 3b): Cinnarizine, a T-type calcium channel blocker and LY97241, a potassium channel inhibitor. L-type calcium channel blockers and anti-inflammatories also showed suggestive evidence of enrichment.

**Table 3.**
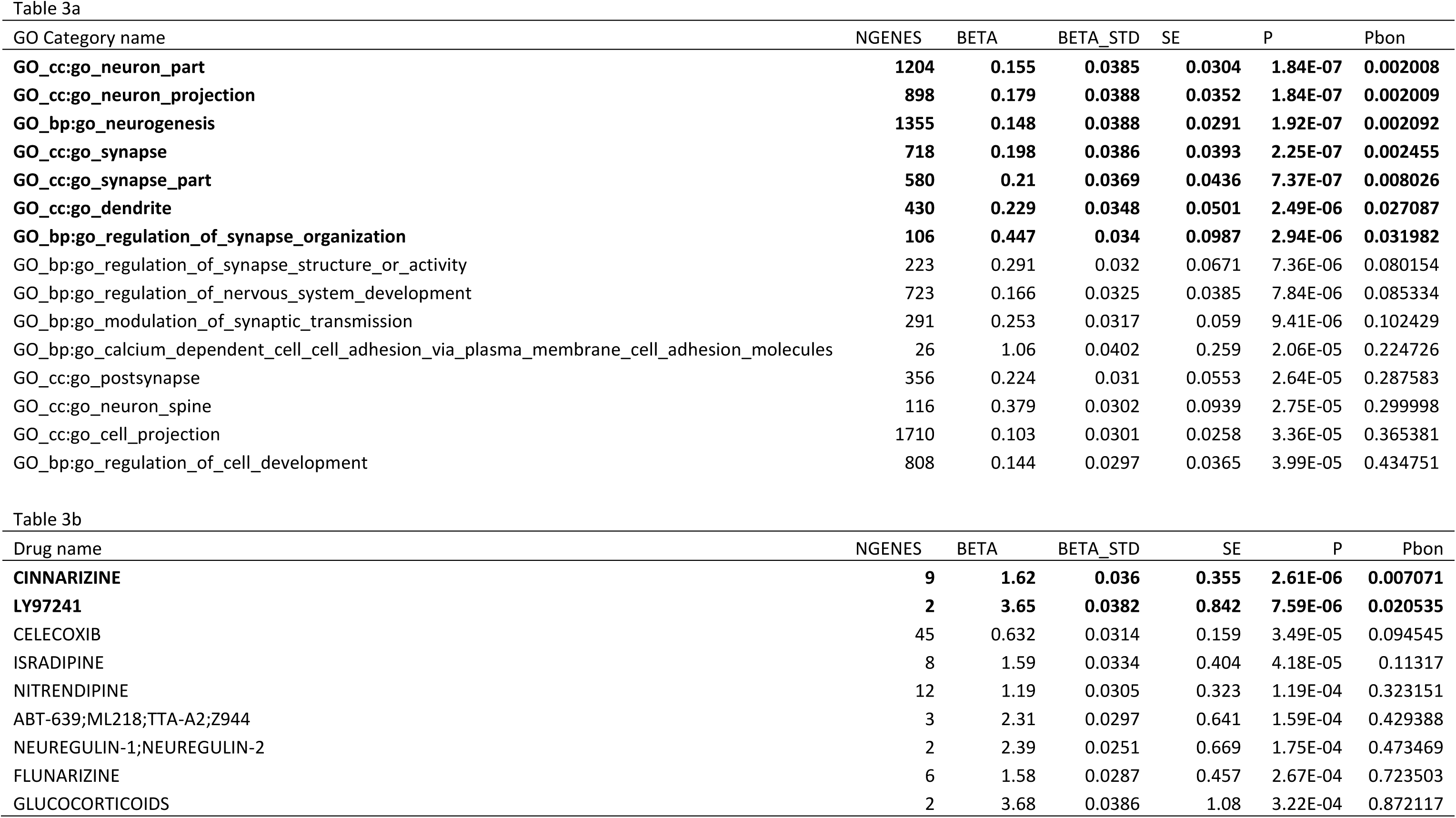

Stratified LD score regression^36^ also demonstrated an enrichment of cell type expression for neuronal tissues only. Notably, genes found in the neuronal expression list of Cahoy^37^ were significantly enriched (p=.0129; Bonferroni-corrected p=.0386), whereas negative results were obtained for genes expressed in oligodendrocytes (p=.4997) and astrocytes (p=.9057). Additionally, using Roadmap annotations, epigenetic enrichment was strongest in fetal brain tissue DNase sites and H3K4me1 primed enhancers; followed by adult cortical H3K27ac active enhancer sites (see Supplementary Table 13 for further details). No enrichment was observed in any non-neuronal tissue.

### Transcriptomic Wide Analysis and Brain Expression Lookups

We performed transcriptome-wide analysis (TWAS, using MetaXcan^38^ with GTEx reference data) on MTAG-derived SNP summary statistics in order to determine whether up- or down-regulation of specific transcripts in specific neural compartments were associated with cognition. (TWAS follows a similar logic to imputation, in that an external reference (in this case, GTEx) is utilized to link available SNP data to tissue-based, gene expression levels.) Several strong transcriptomic associations were specific to individual brain regions such as hippocampus, cortex, or cerebellum. For example, the strongest result in hippocampus was with *DAG1*; TWAS demonstrated that greater expression of this gene in hippocampus was associated with higher cognitive scores. However, this gene was not significantly associated in analyses of other neural tissue types. Similarly, lower levels of *ACTR1A* are associated with cognition, but only in frontal cortex. However, as shown in Supplementary Table 14, most of the strongest TWAS results are tissue non-specific, involving genes such as *AMIGO3, RNF123,* and *RBM6* (Supplementary Figure 11); and a QQ plot revealed that no individual tissue compartment was much more strongly enriched than the others (Supplementary Figure 12). Lookups of GWS SNPs from the MTAG analysis in two brain eQTL databases (BrainEAC^39^ and CommonMind^40^) revealed several additional SNP-eQTL relationships that can explain variance in the cognitive phenotype (Supplementary Tables 15 and 16); the most notable eQTL effect was observed for rs3809912 on chromosome 18. This SNP, which was GWS in the MTAG results (p=7.06E-09), was a strong eQTL for *CEP192* (Bonferroni-corrected p= 3.78E-31 averaged across all neural tissues for expression probe 3779863). This eQTL was confirmed in the CommonMind database (FDR< .01), which demonstrated that expression of 44 independent transcripts in frontal cortex were significantly associated with MTAG SNPs at the FDR< .01 level.

### Genetic Correlations with Other Phenotypes

LD-score regression was carried out across 98 traits in 15 broad phenotypic categories in LD-hub^41^: 1) aging, 2) anthropometric, 3) autoimmune, 4) brain volume, 5) cardiometabolic, 6) education, 7) gycemic, 8) lipids, 9) lung function, 10) neurological, 11) personality, 12) psychiatric, 13) reproductive behavior, 14) sleeping, and 15) smoking behavior. Cognition appeared to be strongly associated with aging, education, personality, neuropsychiatric disorders, reproductive behavior, and smoking behavior. Strong association with parental age at death was observed for both the GWAS and MTAG results. Meanwhile, moderate associations with anthropometric traits were observed, although associations with brain volumes were surprisingly modest, except for total intracranial volume (r_g_ for MTAG results = 0.31). While many of these correlations have been reported previously^12,15,20,25^, two novel results were observed in the present study. First, we report a strong positive genetic correlation between cognitive variables and maternal age at first birth (r_g_ for MTAG results = 0.63, p=2.36E-163) and inverse correlation with parental number of children ever born (r_g_ for MTAG results = -0.2159; p=6.91E-13). It is possible that these effects are mediated by years of higher education, insofar as correlations were even stronger with educational attainment (r_g_ for age at first birth=0.7207, p=2.24E-244; r_g_ for number of children= -0.2623, p=3.34E-18). Second, we observed modest, yet nominally significant, inverse correlations between cognition and autoimmune diseases such as eczema and Crohn’s disease, attaining Bonferroni significance for rheumatoid arthritis (r_g_ for MTAG results = -0.2086; p=1.60E-08); there was also a Bonferroni-significant positive genetic correlation with celiac disease (r_g_ for MTAG results = 0.1922; p=0.0001). While results of cross-trait analyses were largely consistent using either the GWAS results, the MTAG results, or the previously-published educational attainment results, there were notable divergences in correlations with psychiatric phenotypes, especially schizophrenia and bipolar disorder.

## Discussion

Here, we have presented the largest GWAS of cognition to date, with 107,207 individuals phenotypically characterized for performance on standardized tests measuring general cognitive ability. Results were further enhanced by utilizing a novel approach to allow further meta-analysis with a large-scale GWAS of educational attainment, which is highly (though not perfectly) correlated with cognitive ability at the genetic level. With this approach, we were able to identify 70 genomic loci significantly associated with cognition, nearly half of which were novel, and ultimately implicated 350 genes underlying cognitive ability. In total, we found that common SNPs were able to account for nearly half of the overall heritability of the phenotype as determined by prior family studies^42^.

Downstream analysis confirmed an important role for neurodevelopmental processes in cognitive ability, consistent with implications from the GWAS of educational attainment^25^. Significant genes were more strongly enriched for expression in fetal brain tissue than adult tissue; results were also enriched for genes implicated in early neurodevelopmental disorders; and neurogenesis was the most strongly enriched GO biological process. At the same time, it is important to emphasize that adult neural tissues were also strongly represented in the results, and multiple synaptic components were significant in the pathway analysis. In this context, it is noteworthy that many cellular processes necessary for early neurodevelopment are also involved in adult synaptic plasticity. This duality is represented by several significant genes emerging from our analysis: *CELSR3* encodes an atypical cadherin plasma membrane protein involved in long-range axon guidance in neurodevelopment through planar cell polarity signaling^43^, but is also necessary for adult formation of hippocampal glutamatergic synapses^44^. Similarly, *SEMA3F* is a negative regulator of dendritic spine development in adult hippocampus^45^, but embryonically serves as an endogenous chemorepellent, guiding septohippocampal fibers away from non-limbic regions of developing cortex^46^.

To our knowledge, this is the first study of cognition to employ TWAS methodology, which was developed with the hope of isolating expression effects of specific genes within broad GWAS loci. In the present study, a few such genes were isolated, such as *ACTR1A*. This gene, which lies near the GWAS peak at chromosome 10q24, encodes a microtubular dynactin protein involved in retrograde axon transport^47^; other genes at this locus were not significant in the TWAS analysis (although a role in cognition cannot be ruled out, given the limited sample size in the reference brain expression datasets in GTEx). However, most of the genes implicated by TWAS were clustered in a few “hot” genomic loci, which may represent topologically associated domains (TADs) under the control of a shared 3-dimensional chromatin structure^48,49^. Whether effects on cognition are driven by all differentially expressed genes within such loci, or if specific effects can be disentangled through experimental means, remains to be determined.

The overlap of 23 genes from our results with known genes for Mendelian disorders characterized by intellectual disability has several implications. First, this statistically significant enrichment provides partial validation of our MTAG results. Second, genes with known mutations of large effect, when combined with our novel data demonstrating SNPs with smaller regulatory effects on the same phenotype (cognition), can be considered an “allelic series”^50^ – a natural set of experiments powerfully demonstrating directional information (in the form of a dose-response curve) regarding gene function. Such information can be leveraged for the identification of novel drug targets. Third, converging evidence across the Mendelian and GWAS lists can aid interpretation of specific pathways and molecular processes that are necessary to normal neuronal function, and vice versa. For example, two genes on both the Mendelian and GWAS lists (*GMPPB* and *LARGE*) are associated with dystroglycanopathies with mental retardation. This information provides context for the observation that *DAG1*, which encodes dystroglycan 1, is the strongest TWAS result in the hippocampus. *DAG1* is necessary for GABAergic signaling in hippocampal interneurons^50,51^. While dystroglycanopathies are most prominently characterized by muscular dystrophy and retinal abnormalities, it is possible that all of these genes play a role in hippocampal synapse formation that is relevant to normal cognitive ability.

As noted above, one of the most important aims of GWAS studies is the identification of novel drug targets, and the drug set enrichment analysis pointed to potential nootropic mechanisms. Most notably, the strongest signal was for cinnarizine, a T-type calcium channel inhibitor typically prescribed for seasickness. In the present study, we discovered a novel association of cognition to *CACNA1I*, which encodes one component of the voltage-dependent T-Type Cav3.3 channel, and has been previously associated with schizophrenia^52^. While cinnarizine has strong antihistamine activity and may be inappropriate for general cognitive enhancement, a novel agent targeting Cav3.3 has shown nootropic activity in preclinical models. The present study provides supportive evidence for this approach.

It is important to emphasize that uncovering genetic variation underlying general cognitive ability in the healthy population does not have deterministic implications. As has been previously explicated in similar studies, effect sizes for each allele are extremely small (R^2^< 0.1% for even the strongest effects), and the combined effects genome-wide predict only a small proportion (∼2%-4%) of the total variance in hold-out samples (Supplementary Figures 9 and 10). Thus, results of the present study do not hold the potential for individual prediction or classification. Nevertheless, results enhance our understanding of molecular mechanisms underlying cognitive ability, illuminating relationships to other health-relevant traits, and pointing towards specific transcriptomic effects and biological pathways that may form the basis of future cognitive enhancement approaches.

## Methods

### GWAS Quality Control

Markers reported in the prior COGENT study^15^ were updated to build 37 coordinates, but imputed against the HRC reference panel^53^ via the Sanger imputation server. To ensure that markers, allele frequencies, and alleles are aligned to the 1000 genomes phase 3 reference panel^54^, the COGENT summary statistics^15^ were checked using the EasyQC pipeline^55^ which allows summary statistics to be aligned and checked against a reference panel of choice. As both fluid intelligence^20^ and education^25^ summary statistics were imputed to the 1000 genomes phase 3 reference panel, summary statistics were used as provided (URL: https://ctg.cncr.nl/software/summary_statistics; https://www.thessgac.org/data). Further quality control was provided as part of the Multi-trait Analysis for GWAS (MTAG^27^) pipeline. Alleles were aligned against the first dataset, and only SNPs present across datasets were included in the final analysis. The MTAG quality control pipeline is an adaptation of the ‘*munge_sumstats’* function found in LD score regression^39^.

### Fixed Effect Meta-Analysis

Fixed-effect meta-analysis was carried out via METAL^56^. First, fixed-effect meta-analysis was conducted for independent samples reported in the prior COGENT paper^15^ that were not included in the GWAS of fluid intelligence^20^. These cohorts are reported in Supplementary Table 1. The initial meta-analysis resulted in total N = 28,799. Subsequently, further meta-analysis was conducted to combine independent samples from COGENT^15^ and fluid intelligence^20^ resulting in the combined sample size of N = 107,207 for the cognitive performance meta-analysis. Because the fluid intelligence GWAS utilized the sample-size weighted method to perform meta-analysis across its own cohorts, and did not report variance terms, our meta-analysis was conducted using the sample-size weighted method.

### Multi-Trait Analysis for GWAS (MTAG)

To further enrich genetic signals, we employed a newly developed methodology that integrates LD-score genetic regression and meta-analysis techniques across related traits: MTAG^27^. MTAG was applied to the GWAS results applied immediately above, combined with summary statistics from the recent, large-scale educational attainment GWAS^25^. MTAG analysis allows the boosting of genetic signals across related traits, and had been found to be effective in resolving unknown sample overlaps, and generates trait-specific effect estimates weighted by bivariate genetic correlation. The resulting effect estimates and p-values are interpreted in the same manner as single-trait GWAS, which allows standard downstream follow-up analysis on the summary statistics.

### Functional Mapping and Annotation for GWAS

GWAS summary statistics from the METAL meta-analysis and MTAG analysis were entered to the Functional Mapping and Annotation (FUMA) pipeline^28^. The FUMA pipeline enables fast prioritization of genomic variants, genes, and interactive visualization of genomic results with respective to state-of-art bioinformatics resources. Manhattan and QQ plots are produced, and MAGMA gene-based analysis is performed, accounting for gene size and LD structure^32^. Competitive gene set analysis using the Molecular Signature Database (MsigDB 5.2), and brain expression databases from GTEX and BrainEAC were also carried in MAGMA as part of the FUMA pipeline. The pipeline also generates aggregated statistics for independent loci, lead SNP, tagged genes, and supplementary plots – including SNP and Loci annotations. Default clumping parameters are GWAS p-value < 5E-08; r2 threshold to define LD structure of lead SNPs > 0.6; maximum P-value cutoff < 0.05; population for clumping = EUR; Minor Allele Frequency filter > 0.01; maximum distance between LD blocks to merge into a locus < 250kb.

Follow-up queries were then made for top independent loci of the cognitive performance meta-analysis as well as the MTAG results and compared against summary statistics for the prior cognitive^20^ and education^25^ GWAS. For purposes of comparison, loci in which the lead SNPs were within 500kb of each other were considered overlapping.

We compared the list of genes resulting from the MTAG analysis (including all genes within GWS SNP loci, as well as GWS genes identified with MAGMA) with a list of 621 genes known to cause autosomal dominant or autosomal recessive Mendelian disorders featuring intellectual disability; this list is primarily derived from a recent comprehensive review^33^, supplemented by a subsequent large-scale study of consanguineous multiplex families^34^.

FUMA was also utilized to perform competitive gene-set analyses for GO cell compartment and biological process categories. A separate competitive gene-set analysis was also conducted for the drug-based pathways previously described by Gaspar & Breen^35^.

### Polygenic Risk Prediction for independent datasets

To validate that the genetic architecture elucidated via the MTAG methodology, we attempted to predict the phenotypic variance of general cognitive function in two of the independent COGENT cohorts (ASPIS and GCAP). MTAG analysis was conducted as above, but holding out these two cohorts. All polygenic score prediction was conducted using PRSice^57^.

### Stratified LD regression: Cell type Expression and Epigenomics

Functional characterization of GWAS summary statistics were carried out via stratified LD regression to investigate if cognitive heritability is enriched in specific tissue or cell types. Summary statistics were first subjected to baseline partitioned heritability and thereafter passed through cell type functional characterization pipeline ^42^. Cell type characterization includes the DEPICT tissue expression, GTEX tissue expression, IMMGEN immune cell types, CAHOY brain level cell types, and the ROADMAP cell epigenomic marks.

### Transcriptome Wide Analysis and Brain Expression lookups

Downstream transcriptomic wide analysis and brain expression lookups of top SNPs were carried out to examine if specific genomic regions were exclusive associated with specific or multiple brain regions. Transcriptomic wide analysis is carried out via MetaXcan ^43^, which allows for GTEX brain expression mechanisms to be integrated with GWAS summary statistics. Top SNPs obtained from the MTAG GWAS were also subjected to data lookup in the Brain eQTL Almanac (BrainEAC ^44^), as well as CommonMind ^45^ brain expression datasets. GTEX brain tissue expression profiles include the Anterior Cingulate Cortex; Caudate – Basal Ganglia; Cerebellar Hemisphere; Cerebellum; Cortex; Frontal Cortex; Hippocampus; Hypothalamus; Nucleus Accumbens; and Putamen. BrainEAC top SNP lookups were available for the following tissue expression compartments: CRBL: cerebellum; FCTX: frontal cortex; HIPP: Hippocampus; MEDU: medulla; OCTX: occipital cortex; PUTM: putamen; SNIG: substantia nigra; TCTX: temporal cortex; THAL: thalamus; WHMT: white matter; and aveALL: All areas combined. Only one region (the prefrontal cortex) was available for the CommonMind consortium brain expression profile lookup.

### Linkage Disequilibrium Score Regression

LD score regression allows genetic correlations to be computed across traits^58^, which allows further insights to be drawn from understanding the degree to which genetic architecture are shared across traits. To further examine potential traits that overlap with the cognitive architecture from the cognition meta-analysis results and MTAG results, LD score regression was first conducted via the LD-hub pipeline, a centralized trait database^41^. Fifteen broad trait categories were investigated, including: 1) Aging, 2) Anthropometric, 3) Autoimmune, 4) Brain Volume, 5) Cardiometabolic, 6) Education, 7) Glycemic, 8)Lipids, 9) Lung Function, 10) Neurological, 11) Personality, 12) Psychiatric, 13) Reproductive, 14) Sleeping, and 15) Smoking behavior. Very recent results for ADHD^59^ and intracranial volume^60^ were included as additional phenotypes. The same procedures were also carried out for intelligence and education to investigate how the elucidated genetic architecture for the MTAG results differs from prior published works for education and intelligence. Additional LD score regression analysis was conducted between cognitive phenotypes, to examine the degree of genetic architecture overlap across related cognitive traits.

## ACKNOWLEDGMENTS

This work has been supported by grants from the National Institutes of Health (R01 MH079800 and P50 MH080173 to AKM; R01 MH080912 to DCG; K23 MH077807 to KEB; K01 MH085812 to MCK). Data collection for the TOP cohort was supported by the Research Council of Norway, South-East Norway Health Authority, and KG Jebsen Foundation. The NCNG study was supported by Research Council of Norway Grants 154313/V50 and 177458/V50. The NCNG GWAS was financed by grants from the Bergen Research Foundation, the University of Bergen, the Research Council of Norway (FUGE, Psykisk Helse), Helse Vest RHF and Dr Einar Martens Fund. The Helsinki Birth Cohort Study has been supported by grants from the Academy of Finland, the Finnish Diabetes Research Society, Folkhälsan Research Foundation, Novo Nordisk Foundation, Finska Läkaresällskapet, Signe and Ane Gyllenberg Foundation, University of Helsinki, Ministry of Education, Ahokas Foundation, Emil Aaltonen Foundation. For the LBC1936 cohort, phenotype collection was supported by The Disconnected Mind project. Genotyping was funded by the UK Biotechnology and Biological Sciences Research Council (BBSRC grant No. BB/F019394/1). The work was undertaken by The University of Edinburgh Centre for Cognitive Ageing and Cognitive Epidemiology, part of the cross council Lifelong Health and Wellbeing Initiative, which is funded by the Medical Research Council and the Biotechnology and Biological Sciences Research Council (MR/K026992/1). The CAMH work was supported by the CAMH Foundation and the Canadian Institutes of Health Research. The Duke Cognition Cohort (DCC) acknowledges K. Linney, J.M. McEvoy, P. Hunt, V. Dixon, T. Pennuto, K. Cornett, D. Swilling, L. Phillips, M. Silver, J. Covington, N. Walley, J. Dawson, H. Onabanjo, P. Nicoletti, A. Wagoner, J. Elmore, L. Bevan, J. Hunkin and R. Wilson for recruitment and testing of subjects. DCC also acknowledges the Ellison Medical Foundation New Scholar award AG-NS-0441-08 for partial funding of this study as well as the National Institute of Mental Health of the National Institutes of Health under award number K01MH098126. The UCLA Consortium for Neuropsychiatric Phenomics (CNP) study acknowledges the following sources of funding from the NIH: Grants UL1DE019580 and PL1MH083271 (RMB), RL1MH083269 (TDC), RL1DA024853 (EL) and PL1NS062410. The ASPIS study was supported by National Institute of Mental Health research grants R01MH085018 and R01MH092515 to Dr. Dimitrios Avramopoulos. Support for the Duke Neurogenetics Study was provided the National Institutes of Health (R01 DA033369 and R01 AG049789 to ARH) and by a National Science Foundation Graduate Research Fellowship to MAS. Recruitment, genotyping and analysis of the TCD healthy control samples were supported by Science Foundation Ireland (grants 12/IP/1670, 12/IP/1359 and 08/IN.1/B1916).

Data access for several cohorts used in this study was provided by the National Center for Biotechnology Information (NCBI) database of Genotypes and Phenotypes (dbGaP). dbGaP accession numbers for these cohorts were:

Cardiovascular Health Study (CHS): phs000287.v4.p1, phs000377.v5.p1, and phs000226.v3.p1 Framingham Heart Study (FHS): phs000007.v23.p8 and phs000342.v11.p8

Multi-Site Collaborative Study for Genotype-Phenotype Associations in Alzheimer’s Disease (GENADA): phs000219.v1.p1

Long Life Family Study (LLFS): phs000397.v1.p1

Genetics of Late Onset Alzheimer’s Disease Study (LOAD): phs000168.v1.p1

Minnesota Center for Twin and Family Research (MCTFR): phs000620.v1.p1

Philadelphia Neurodevelopmental Cohort (PNC): phs000607.v1.p1

The acknowledgment statements for these cohorts are found below:

Framingham Heart Study: The Framingham Heart Study is conducted and supported by the National Heart, Lung, and Blood Institute (NHLBI) in collaboration with Boston University (Contract No. N01-HC-25195 and HHSN268201500001I). This manuscript was not prepared in collaboration with investigators of the Framingham Heart Study and does not necessarily reflect the opinions or views of the Framingham Heart Study, Boston University, or NHLBI. Funding for SHARe Affymetrix genotyping was provided by NHLBI Contract N02-HL-64278. SHARe Illumina genotyping was provided under an agreement between Illumina and Boston University.

Cardiovascular Health Study: This research was supported by contracts HHSN268201200036C, HHSN268200800007C, N01-HC-85079, N01-HC-85080, N01-HC-85081, N01-HC-85082, N01-HC-85083, N01-HC-85084, N01-HC-85085, N01-HC-85086, N01-HC-35129, N01 HC-15103, N01 HC-55222, N01-HC-75150, N01-HC-45133, and N01-HC-85239; grant numbers U01 HL080295 and U01 HL130014 from the National Heart, Lung, and Blood Institute, and R01 AG-023629 from the National Institute on Aging, with additional contribution from the National Institute of Neurological Disorders and Stroke. A full list of principal CHS investigators and institutions can be found at https://chs-nhlbi.org/pi. This manuscript was not prepared in collaboration with CHS investigators and does not necessarily reflect the opinions or views of CHS, or the NHLBI. Support for the genotyping through the CARe Study was provided by NHLBI Contract N01-HC-65226. Support for the Cardiovascular Health Study Whole Genome Study was provided by NHLBI grant HL087652. Additional support for infrastructure was provided by HL105756 and additional genotyping among the African-American cohort was supported in part by HL085251, DNA handling and genotyping at Cedars-Sinai Medical Center was supported in part by National Center for Research Resources grant UL1RR033176, now at the National Center for Advancing Translational Technologies CTSI grant UL1TR000124; in addition to the National Institute of Diabetes and Digestive and Kidney Diseases grant DK063491 to the Southern California Diabetes Endocrinology Research Center.

Multi-Site Collaborative Study for Genotype-Phenotype Associations in Alzheimer’s Disease: The genotypic and associated phenotypic data used in the study were provided by the GlaxoSmithKline, R&D Limited. Details on data acquisition have been published previously in: Li H, Wetten S, Li L, St Jean PL, Upmanyu R, Surh L, Hosford D, Barnes MR, Briley JD, Borrie M, Coletta N, Delisle R, Dhalla D, Ehm MG, Feldman HH, Fornazzari L, Gauthier S, Goodgame N, Guzman D, Hammond S, Hollingworth P, Hsiung GY, Johnson J, Kelly DD, Keren R, Kertesz A, King KS, Lovestone S, Loy-English I, Matthews PM, Owen MJ, Plumpton M, Pryse-Phillips W, Prinjha RK, Richardson JC, Saunders A, Slater AJ, St George-Hyslop PH, Stinnett SW, Swartz JE, Taylor RL, Wherrett J, Williams J, Yarnall DP, Gibson RA, Irizarry MC, Middleton LT, Roses AD. Candidate single-nucleotide polymorphisms from a genomewide association study of Alzheimer disease. Arch Neurol., Jan;65(1):45-53, 2008 (PMID: 17998437).

Filippini N, Rao A, Wetten S, Gibson RA, Borrie M, Guzman D, Kertesz A, Loy-English I, Williams J, Nichols T, Whitcher B, Matthews PM. Anatomically-distinct genetic associations of APOE epsilon4 allele load with regional cortical atrophy in Alzheimer’s disease. Neuroimage, Feb 1;44(3):724-8, 2009. (PMID: 19013250).

Genetics of Late Onset Alzheimer’s Disease Study: Funding support for the “Genetic Consortium for Late Onset Alzheimer’s Disease” was provided through the Division of Neuroscience, NIA. The Genetic Consortium for Late Onset Alzheimer’s Disease includes a genome-wide association study funded as part of the Division of Neuroscience, NIA. Assistance with phenotype harmonization and genotype cleaning, as well as with general study coordination, was provided by Genetic Consortium for Late Onset Alzheimer’s Disease. A list of contributing investigators is available at https://www.ncbi.nlm.nih.gov/projects/gap/cgi-bin/study.cgi?study_id=phs000168.v1.p1

Long Life Family Study: Funding support for the Long Life Family Study was provided by the Division of Geriatrics and Clinical Gerontology, National Institute on Aging. The Long Life Family Study includes GWAS analyses for factors that contribute to long and healthy life. Assistance with phenotype harmonization and genotype cleaning as well as with general study coordination, was provided by the Division of Geriatrics and Clinical Gerontology, National Institute on Aging. Support for the collection of datasets and samples were provided by Multicenter Cooperative Agreement support by the Division of Geriatrics and Clinical Gerontology, National Institute on Aging (UO1AG023746; UO1023755; UO1023749; UO1023744; UO1023712). Funding support for the genotyping which was performed at the Johns Hopkins University Center for Inherited Disease Research was provided by the National Institute on Aging, National Institutes of Health.

Minnesota Center for Twin and Family Research: This project was led by William G. Iacono, PhD. And Matthew K. McGue, PhD (Co-Principal Investigators) at the University of Minnesota, Minneapolis, MN, USA. Co-investigators from the same institution included: Irene J. Elkins, Margaret A. Keyes, Lisa N. Legrand, Stephen M. Malone, William S. Oetting, Michael B. Miller, and Saonli Basu. Funding support for this project was provided through NIDA (U01 DA 024417). Other support for sample ascertainment and data collection came from several grants: R37 DA 05147, R01 AA 09367, R01 AA 11886, R01 DA 13240, R01 MH 66140.

Philadelphia Neurodevelopmental Cohort: Support for the collection of the data sets was provided by grant RC2MH089983 awarded to Raquel Gur, MD, and RC2MH089924 awarded to Hakon Hakonarson, MD, PhD. All subjects were recruited through the Center for Applied Genomics at The Children’s Hospital in Philadelphia.

## References

1. Wood AR, Esko T, Yang J, Vedantam S, Pers TH, Gustafsson S et al. Defining the role of common variation in the genomic and biological architecture of adult human height. Nat Genet 2014; 46: 1173–1186.

2. Locke AE, Kahali B, Berndt SI, Justice AE, Pers TH, Day FR et al. Genetic studies of body mass index yield new insights for obesity biology. Nature 2015; 518: 197–206.

3. Deary IJ, Johnson W, Houlihan LM. Genetic foundations of human intelligence. Hum Genet 2009; 126: 215–32.

4. Davies G, Tenesa A, Payton A, Yang J, Harris SE, Liewald D et al. Genome-wide association studies establish that human intelligence is highly heritable and polygenic. Mol Psychiatry 2011; 16: 996–1005

5. Keefe RS, Harvey PD. Cognitive impairment in schizophrenia. Handb Exp Pharmacol 2012; 213:11–37.

6. Burdick KE, Goldberg TE, Cornblatt B a, Keefe RS, Gopin CB, Derosse P et al. The MATRICS consensus cognitive battery in patients with bipolar I disorder. Neuropsychopharmacology 2011; 36: 1587–1592.

7. Ferreri F, Lapp LK, Peretti C-S. Current research on cognitive aspects of anxiety disorders. Curr Opin Psychiatry 2011; 24: 49–54.

8. Snyder HR. Major depressive disorder is associated with broad impairments on neuropsychological measures of executive function: a meta-analysis and review. Psychol Bull 2013; 139: 81–132

9. Lencz, T. et al. Molecular genetic evidence for overlap between general cognitive ability and risk for schizophrenia: a report from the Cognitive Genomics consorTium (COGENT). Mol. Psychiatry 19, 168–174 (2014).

10. Smeland OB, Frei O, Kauppi K, Hill WD, Li W, Wang Y, Krull F, Bettella F, Eriksen JA, Witoelar A, Davies G, Fan CC, Thompson WK, Lam M, Lencz T, Chen CH, Ueland T, Jönsson EG, Djurovic S, Deary IJ, Dale AM, Andreassen OA; NeuroCHARGE (Cohorts for Heart and Aging Research in Genomic Epidemiology) Cognitive Working Group. Identification of Genetic Loci Jointly Influencing Schizophrenia Risk and the Cognitive Traits of Verbal-Numerical Reasoning, Reaction Time, and General Cognitive Function. JAMA Psychiatry. 2017 Jul 26. [Epub ahead of print]

11. Stergiakouli E, Martin J, Hamshere ML, Heron J, St Pourcain B, Timpson NJ, Thapar A, Davey Smith G. Association between polygenic risk scores for attention-deficit hyperactivity disorder and educational and cognitive outcomes in the general population. Int J Epidemiol. 2017 Apr 1;46(2):421–428.

12. Hagenaars SP, Harris SE, Davies G, Hill WD, Liewald DC, Ritchie SJ, Marioni RE, Fawns-Ritchie C, Cullen B, Malik R; METASTROKE Consortium, International Consortium for Blood Pressure GWAS; SpiroMeta Consortium; CHARGE Consortium Pulmonary Group, CHARGE Consortium Aging and Longevity Group, Worrall BB, Sudlow CL, Wardlaw JM, Gallacher J, Pell J, McIntosh AM, Smith DJ, Gale CR, Deary IJ. Shared genetic aetiology between cognitive functions and physical and mental health in UK Biobank (N=112 151) and 24 GWAS consortia. Mol Psychiatry. 2016 Nov;21(11):1624–1632.

13. Johnson W, te Nijenhuis J, Bouchard Jr TJ. Still just 1 g: Consistent results from five test batteries. Intelligence 2008; 36: 81–95.

14. Benyamin B, Pourcain B, Davis OS, Davies G, Hansell NK, Brion M-J a et al. Childhood intelligence is heritable, highly polygenic and associated with FNBP1L. Mol Psychiatry 2014; 19: 253–8.

15. Trampush, J. W. et al. GWAS meta-analysis reveals novel loci and genetic correlates for general cognitive function: a report from the COGENT consortium. Mol. Psychiatry 22, 336–345 (2017).

16. Davies G, Armstrong N, Bis JC, Bressler J, Chouraki V, Giddaluru S et al. Genetic contributions to variation in general cognitive function: a meta-analysis of genome-wide association studies in the CHARGE consortium (N=53 949). Mol Psychiatry 2015: 1–10.

17. Weedon MN, Lango H, Lindgren CM, Wallace C, Evans DM, Mangino M, Freathy RM, Perry JR, Stevens S, Hall AS, Samani NJ, Shields B, Prokopenko I, Farrall M, Dominiczak A; Diabetes Genetics Initiative; Wellcome Trust Case Control Consortium, Johnson T, Bergmann S, Beckmann JS, Vollenweider P, Waterworth DM, Mooser V, Palmer CN, Morris AD, Ouwehand WH; Cambridge GEM Consortium, Zhao JH, Li S, Loos RJ, Barroso I, Deloukas P, Sandhu MS, Wheeler E, Soranzo N, Inouye M, Wareham NJ, Caulfield M, Munroe PB, Hattersley AT, McCarthy MI, Frayling TM. Genome-wide association analysis identifies 20 loci that influence adult height. Nat Genet. 2008 May;40(5):575–83.

18. Gudbjartsson DF, Walters GB, Thorleifsson G, Stefansson H, Halldorsson BV, Zusmanovich P, Sulem P, Thorlacius S, Gylfason A, Steinberg S, Helgadottir A, Ingason A, Steinthorsdottir V, Olafsdottir EJ, Olafsdottir GH, Jonsson T, Borch-Johnsen K, Hansen T, Andersen G, Jorgensen T, Pedersen O, Aben KK, Witjes JA, Swinkels DW, den Heijer M, Franke B, Verbeek AL, Becker DM, Yanek LR, Becker LC, Tryggvadottir L, Rafnar T, Gulcher J, Kiemeney LA, Kong A, Thorsteinsdottir U, Stefansson K. Many sequence variants affecting diversity of adult human height. Nat Genet. 2008 May;40(5):609–15.

19. Trampush JW, Lencz T, Knowles E, Davies G, Guha S, Pe’er I et al. Independent evidence for an association between general cognitive ability and a genetic locus for educational attainment. Am J Med Genet Part B Neuropsychiatr Genet 2015; 168B:363–373.

20. Sniekers, S. et al. Genome-wide association meta-analysis of 78,308 individuals identifies new loci and genes influencing human intelligence. Nat. Genet. 49, 1107–1112 (2017).

21. Rietveld CA, Esko T, Davies G, Pers TH, Turley P, Benyamin B, Chabris CF, Emilsson V, Johnson AD, Lee JJ, de Leeuw C, Marioni RE, Medland SE, Miller MB, Rostapshova O, van der Lee SJ, Vinkhuyzen AA, Amin N, Conley D, Derringer J, van Duijn CM, Fehrmann R, Franke L, Glaeser EL, Hansell NK, Hayward C, Iacono WG, Ibrahim-Verbaas C, Jaddoe V, Karjalainen J, Laibson D, Lichtenstein P, Liewald DC, Magnusson PK, Martin NG, McGue M, McMahon G, Pedersen NL, Pinker S, Porteous DJ, Posthuma D, Rivadeneira F, Smith BH, Starr JM, Tiemeier H, Timpson NJ, Trzaskowski M, Uitterlinden AG, Verhulst FC, Ward ME, Wright MJ, Davey Smith G, Deary IJ, Johannesson M, Plomin R, Visscher PM, Benjamin DJ, Cesarini D, Koellinger PD. Common genetic variants associated with cognitive performance identified using the proxy-phenotype method. Proc Natl Acad Sci U S A. 2014 Sep 23;111(38):13790–4.

22. Belsky DW, Moffitt TE, Corcoran DL, Domingue B, Harrington H, Hogan S, Houts R, Ramrakha S, Sugden K, Williams BS, Poulton R, Caspi A. The Genetics of Success: How Single-Nucleotide Polymorphisms Associated With Educational Attainment Relate to Life-Course Development. Psychol Sci. 2016 Jul;27(7):957–72.

23. Johnson W, Deary IJ, Silventoinen K, Tynelius P, Rasmussen F. Family background buys an education in Minnesota but not in Sweden. Psychol Sci. 2010 Sep;21(9):1266–73.

24. Rietveld CA, Medland SE, Derringer J, Yang J, Esko T, Martin NW, Westra HJ, Shakhbazov K, Abdellaoui A, Agrawal A, Albrecht E, Alizadeh BZ, Amin N, Barnard J, Baumeister SE, Benke KS, Bielak LF, Boatman JA, Boyle PA, Davies G, de Leeuw C, Eklund N, Evans DS, Ferhmann R, Fischer K, Gieger C, Gjessing HK, Hägg S, Harris JR, Hayward C, Holzapfel C, Ibrahim-Verbaas CA, Ingelsson E, Jacobsson B, Joshi PK, Jugessur A, Kaakinen M, Kanoni S, Karjalainen J, Kolcic I, Kristiansson K, Kutalik Z, Lahti J, Lee SH, Lin P, Lind PA, Liu Y, Lohman K, Loitfelder M, McMahon G, Vidal PM, Meirelles O, Milani L, Myhre R, Nuotio ML, Oldmeadow CJ, Petrovic KE, Peyrot WJ, Polasek O, Quaye L, Reinmaa E, Rice JP, Rizzi TS, Schmidt H, Schmidt R, Smith AV, Smith JA, Tanaka T, Terracciano A, van der Loos MJ, Vitart V, Völzke H, Wellmann J, Yu L, Zhao W, Allik J, Attia JR, Bandinelli S, Bastardot F, Beauchamp J, Bennett DA, Berger K, Bierut LJ, Boomsma DI, Bültmann U, Campbell H, Chabris CF, Cherkas L, Chung MK, Cucca F, de Andrade M, De Jager PL, De Neve JE, Deary IJ, Dedoussis GV, Deloukas P, Dimitriou M, Eiríksdóttir G, Elderson MF, Eriksson JG, Evans DM, Faul JD, Ferrucci L, Garcia ME, Grönberg H, Guðnason V, Hall P, Harris JM, Harris TB, Hastie ND, Heath AC, Hernandez DG, Hoffmann W, Hofman A, Holle R, Holliday EG, Hottenga JJ, Iacono WG, Illig T, Järvelin MR, Kähönen M, Kaprio J, Kirkpatrick RM, Kowgier M, Latvala A, Launer LJ, Lawlor DA, Lehtimäki T, Li J, Lichtenstein P, Lichtner P, Liewald DC, Madden PA, Magnusson PK, Mäkinen TE, Masala M, McGue M, Metspalu A, Mielck A, Miller MB, Montgomery GW, Mukherjee S, Nyholt DR, Oostra BA, Palmer LJ, Palotie A, Penninx BW, Perola M, Peyser PA, Preisig M, Räikkönen K, Raitakari OT, Realo A, Ring SM, Ripatti S, Rivadeneira F, Rudan I, Rustichini A, Salomaa V, Sarin AP, Schlessinger D, Scott RJ, Snieder H, St Pourcain B, Starr JM, Sul JH, Surakka I, Svento R, Teumer A; LifeLines Cohort Study, Tiemeier H, van Rooij FJ, Van Wagoner DR, Vartiainen E, Viikari J, Vollenweider P, Vonk JM, Waeber G, Weir DR, Wichmann HE, Widen E, Willemsen G, Wilson JF, Wright AF, Conley D, Davey-Smith G, Franke L, Groenen PJ, Hofman A, Johannesson M, Kardia SL, Krueger RF, Laibson D, Martin NG, Meyer MN, Posthuma D, Thurik AR, Timpson NJ, Uitterlinden AG, van Duijn CM, Visscher PM, Benjamin DJ, Cesarini D, Koellinger PD. GWAS of 126,559 individuals identifies genetic variants associated with educational attainment. Science. 2013 Jun 21;340(6139):1467–71.

25. Okbay A, Beauchamp JP, Fontana MA, Lee JJ, Pers TH, Rietveld CA, Turley P, Chen GB, Emilsson V, Meddens SF, Oskarsson S, Pickrell JK, Thom K, Timshel P, de Vlaming R, Abdellaoui A, Ahluwalia TS, Bacelis J, Baumbach C, Bjornsdottir G, Brandsma JH, Pina Concas M, Derringer J, Furlotte NA, Galesloot TE, Girotto G, Gupta R, Hall LM, Harris SE, Hofer E, Horikoshi M, Huffman JE, Kaasik K, Kalafati IP, Karlsson R, Kong A, Lahti J, van der Lee SJ, deLeeuw C, Lind PA, Lindgren KO, Liu T, Mangino M, Marten J, Mihailov E, Miller MB, van der Most PJ, Oldmeadow C, Payton A, Pervjakova N, Peyrot WJ, Qian Y, Raitakari O, Rueedi R, Salvi E, Schmidt B, Schraut KE, Shi J, Smith AV, Poot RA, St Pourcain B, Teumer A, Thorleifsson G, Verweij N, Vuckovic D, Wellmann J, Westra HJ, Yang J, Zhao W, Zhu Z, Alizadeh BZ, Amin N, Bakshi A, Baumeister SE, Biino G, Bønnelykke K, Boyle PA, Campbell H, Cappuccio FP, Davies G, De Neve JE, Deloukas P, Demuth I, Ding J, Eibich P, Eisele L, Eklund N, Evans DM, Faul JD, Feitosa MF, Forstner AJ, Gandin I, Gunnarsson B, Halldórsson BV, Harris TB, Heath AC, Hocking LJ, Holliday EG, Homuth G, Horan MA, Hottenga JJ, de Jager PL, Joshi PK, Jugessur A, Kaakinen MA, Kähönen M, Kanoni S, Keltigangas-Järvinen L, Kiemeney LA, Kolcic I, Koskinen S, Kraja AT, Kroh M, Kutalik Z, Latvala A, Launer LJ, Lebreton MP, Levinson DF, Lichtenstein P, Lichtner P, Liewald DC; LifeLines Cohort Study, Loukola A, Madden PA, Mägi R, Mäki-Opas T, Marioni RE, Marques-Vidal P, Meddens GA, McMahon G, Meisinger C, Meitinger T, Milaneschi Y, Milani L, Montgomery GW, Myhre R, Nelson CP, Nyholt DR, Ollier WE, Palotie A, Paternoster L, Pedersen NL, Petrovic KE, Porteous DJ, Räikkönen K, Ring SM, Robino A, Rostapshova O, Rudan I, Rustichini A, Salomaa V, Sanders AR, Sarin AP, Schmidt H, Scott RJ, Smith BH, Smith JA, Staessen JA, Steinhagen-Thiessen E, Strauch K, Terracciano A, Tobin MD, Ulivi S, Vaccargiu S, Quaye L, van Rooij FJ, Venturini C, Vinkhuyzen AA, Völker U, Völzke H, Vonk JM, Vozzi D, Waage J, Ware EB, Willemsen G, Attia JR, Bennett DA, Berger K, Bertram L, Bisgaard H, Boomsma DI, Borecki IB, Bültmann U, Chabris CF, Cucca F, Cusi D, Deary IJ, Dedoussis GV, van Duijn CM, Eriksson JG, Franke B, Franke L, Gasparini P, Gejman PV, Gieger C, Grabe HJ, Gratten J, Groenen PJ, Gudnason V, van der Harst P, Hayward C, Hinds DA, Hoffmann W, Hyppönen E, Iacono WG, Jacobsson B, Järvelin MR, Jöckel KH, Kaprio J, Kardia SL, Lehtimäki T, Lehrer SF, Magnusson PK, Martin NG, McGue M, Metspalu A, Pendleton N, Penninx BW, Perola M, Pirastu N, Pirastu M, Polasek O, Posthuma D, Power C, Province MA, Samani NJ, Schlessinger D, Schmidt R, Sørensen TI, Spector TD, Stefansson K, Thorsteinsdottir U, Thurik AR, Timpson NJ, Tiemeier H, Tung JY, Uitterlinden AG, Vitart V, Vollenweider P, Weir DR, Wilson JF, Wright AF, Conley DC, Krueger RF, Davey Smith G, Hofman A, Laibson DI, Medland SE, Meyer MN, Yang J, Johannesson M, Visscher PM, Esko T, Koellinger PD, Cesarini D, Benjamin DJ. Genome-wide association study identifies 74 loci associated with educational attainment. Nature. 2016 May 26;533(7604):539–42

26. Davies, G. et al. Genome-wide association study of cognitive functions and educational attainment in UK Biobank (N = 112 151). Mol. Psychiatry (2016).

27. Turley P, Walters RK, Maghzian O, Okbay A, Lee JJ, Fontana MA, Nguyen-Viet TA, Wedow R, Zacher M, Furlotte NA, 23andMe Research Team, Social Science Genetic Association Consortium, Magnusson P, Oskarsson S, Johannesson M, Visscher PM, Laibson D, Cesarini D, Neale B, Benjamin DJ. MTAG: Multi-Trait Analysis of GWAS.

28. Watanabe, K., Taskesen, E., van Bochoven, A. & Posthuma, D. FUMA: Functional mapping and annotation of genetic associations. bioRxiv (2017). doi:10.1101/110023

29. Guha S, Rees E, Darvasi A, Ivanov D, Ikeda M, Bergen SE, Magnusson PK, Cormican P, Morris D, Gill M, Cichon S, Rosenfeld JA, Lee A, Gregersen PK, Kane JM, Malhotra AK, Rietschel M, Nöthen MM, Degenhardt F, Priebe L, Breuer R, Strohmaier J, Ruderfer DM, Moran JL, Chambert KD, Sanders AR, Shi J, Kendler K, Riley B, O’Neill T, Walsh D, Malhotra D, Corvin A, Purcell S, Sklar P, Iwata N, Hultman CM, Sullivan PF, Sebat J, McCarthy S, Gejman PV, Levinson DF, Owen MJ, O’Donovan MC, Lencz T, Kirov G; Molecular Genetics of Schizophrenia Consortium; Wellcome Trust Case Control Consortium 2. Implication of a rare deletion at distal 16p11.2 in schizophrenia. JAMA Psychiatry. 2013;70(3):253–60.

30. Myers AJ, Kaleem M, Marlowe L, Pittman AM, Lees AJ, Fung HC et al. The H1c haplotype at the MAPT locus is associated with Alzheimer’s disease. Hum Mol Genet 2005; 14: 2399–404

31. Cooper GM, Coe BP, Girirajan S, Rosenfeld JA, Vu TH, Baker C et al. A copy number variation morbidity map of developmental delay. Nat Genet 2011; 43: 838–46.

32. de Leeuw, C. A., Mooij, J. M., Heskes, T. & Posthuma, D. MAGMA: Generalized Gene-Set Analysis of GWAS Data. PLoS Comput. Biol. 11, (2015).

33. Vissers LE, Gilissen C, Veltman JA. Genetic studies in intellectual disability and related disorders. Nat Rev Genet. 2016 Jan;17(1):9–18.

34. Harripaul R, Vasli N, Mikhailov A, Rafiq MA, Mittal K, Windpassinger C, Sheikh TI, Noor A, Mahmood H, Downey S, Johnson M, Vleuten K, Bell L, Ilyas M, Khan FS, Khan V, Moradi M, Ayaz M, Naeem F, Heidari A, Ahmed I, Ghadami S, Agha Z, Zeinali S, Qamar R, Mozhdehipanah H, John P, Mir A, Ansar M, French L, Ayub M, Vincent JB. Mapping autosomal recessive intellectual disability: combined microarray and exome sequencing identifies 26 novel candidate genes in 192 consanguineous families. Mol Psychiatry. 2017 Apr 11. [Epub ahead of print]

35. Gaspar, H. A. & Breen, G. Pathways analyses of schizophrenia GWAS focusing on known and novel drug targets. bioRxiv (2016). doi:10.1101/091264

36. Finucane, H. et al. Heritability enrichment of specifically expressed genes identifies disease-relevant tissues and cell types. bioRxiv (2017). doi:10.1101/103069

37. Cahoy JD, Emery B, Kaushal A, Foo LC, Zamanian JL, Christopherson KS, Xing Y, Lubischer JL, Krieg PA, Krupenko SA, Thompson WJ, Barres BA. A transcriptome database for astrocytes, neurons, and oligodendrocytes: a new resource for understanding brain development and function. J Neurosci. 2008 Jan 2;28(1):264–78

38. Barbeira, A. et al. MetaXcan: Summary Statistics Based Gene-Level Association Method Infers Accurate PrediXcan Results. bioRxiv (2016). doi:10.1101/045260

39. Ramasamy, A. et al. Genetic variability in the regulation of gene expression in ten regions of the human brain. Nat Neurosci 17, 1418–1428 (2014).

40. Hauberg ME, Zhang W, Giambartolomei C, Franzén O, Morris DL, Vyse TJ, Ruusalepp A; CommonMind Consortium, Sklar P, Schadt EE, Björkegren JLM, Roussos P. Large-Scale Identification of Common Trait and Disease Variants Affecting Gene Expression. Am J Hum Genet. 2017 Jun 1;100(6):885–894.

41. Zheng, J. et al. LD Hub: a centralized database and web interface to perform LD score regression that maximizes the potential of summary level GWAS data for SNP heritability and genetic correlation analysis. Bioinformatics 33, 272–279 (2017)

42. Plomin R, Deary IJ. Genetics and intelligence differences: five special findings. Mol Psychiatry. 2015 Feb;20(1):98–108.

42. McCarthy, S. et al. A reference panel of 64,976 haplotypes for genotype imputation. Nat Genet 48, 1279–1283 (2016).

43. Chai G, Goffinet AM, Tissir F. Celsr3 and Fzd3 in axon guidance. Int J Biochem Cell Biol. 2015 Jul;64:11–4.

43. The 1000 Genomes Project Consortium. A global reference for human genetic variation. Nature 526, 68–74 (2015).

44. Thakar S, Wang L, Yu T, Ye M, Onishi K, Scott J, Qi J, Fernandes C, Han X, Yates JR 3rd, Berg DK, Zou Y. Evidence for opposing roles of Celsr3 and Vangl2 in glutamatergic synapse formation. Proc Natl Acad Sci U S A. 2017 Jan 24;

44. Winkler, T. W. et al. Quality control and conduct of genome-wide association meta-analyses. Nat. Protoc. 9, 1192–1212 (2014).

45. Tran TS, Rubio ME, Clem RL, Johnson D, Case L, Tessier-Lavigne M, Huganir RL, Ginty DD, Kolodkin AL. Secreted semaphorins control spine distribution and morphogenesis in the postnatal CNS Nature. 2009 Dec 24;462(7276):1065–9.

46. Pascual M, Pozas E, Soriano E. Role of class 3 semaphorins in the development and maturation of the septohippocampal pathway. Hippocampus. 2005;15(2):184–202.

47. Moughamian AJ1, Osborn GE, Lazarus JE, Maday S, Holzbaur EL. Ordered recruitment of dynactin to the microtubule plus-end is required for efficient initiation of retrograde axonal transport. J Neurosci. 2013 Aug 7;33(32):13190–203. doi:10.1523/JNEUROSCI.0935-13.2013.

48. Grubert F, Zaugg JB, Kasowski M, Ursu O, Spacek DV, Martin AR, Greenside P, Srivas R, Phanstiel DH, Pekowska A, Heidari N, Euskirchen G, Huber W, Pritchard JK, Bustamante CD, Steinmetz LM, Kundaje A, Snyder M. Genetic Control of Chromatin States in Humans Involves Local and Distal Chromosomal Interactions. Cell. 2015 Aug 27;162(5):1051–65.

49. Gonzalez-Sandoval A, Gasser SM. On TADs and LADs: Spatial Control Over Gene Expression. Trends Genet. 2016 Aug;32(8):485–95. doi: 10.1016/j.tig.2016.05.004. Epub 2016 Jun 13.

50. Plenge RM, Scolnick EM, Altshuler D. Validating therapeutic targets through human genetics. Nat Rev Drug Discov. 2013 Aug;12(8):581–94.

51. Früh S, Romanos J, Panzanelli P, Bürgisser D, Tyagarajan SK, Campbell KP, Santello M, Fritschy JM. Neuronal Dystroglycan Is Necessary for Formation and Maintenance of Functional CCK-Positive Basket Cell Terminals on Pyramidal Cells. J Neurosci. 2016 Oct 5;36(40):10296–10313.

52. Schizophrenia Working Group of the Psychiatric Genomics Consortium. Biological insights from 108 schizophrenia-associated genetic loci. Nature. 2014 Jul 24;511(7510):421–7.

53. Moriguchi S, Shioda N, Yamamoto Y, Tagashira H, Fukunaga K. The T-type voltage-gated calcium channel as a molecular target of the novel cognitive enhancer ST101: enhancement of long-term potentiation and CaMKII autophosphorylation in rat cortical slices. J Neurochem. 2012 Apr;121(1):44–53.

54. Willer, C. J., Li, Y. & Abecasis, G. R. METAL: fast and efficient meta-analysis of genome-wide association scans. Bioinforma. Oxf. Engl. 26, 2190–2191 (2010).

55. Euesden, J., Lewis, C. M. & O’Reilly, P. F. PRSice: Polygenic Risk Score software. Bioinforma. Oxf. Engl. 31, 1466–1468 (2015).

56. Bulik-Sullivan, B. et al. An Atlas of Genetic Correlations across Human Diseases and Traits. bioRxiv 014498 (2015). doi:10.1101/014498

57. Demontis, D. et al. Discovery Of The First Genome-Wide Significant Risk Loci For ADHD. bioRxiv (2017). doi:10.1101/145581

58. Adams, H. H. H. et al. Novel genetic loci underlying human intracranial volume identified through genome-wide association. Nat. Neurosci. 19, 1569–1582 (2016).

